# Novel pre-clinical model for CDKL5 Deficiency Disorder

**DOI:** 10.1101/2021.04.07.438335

**Authors:** Rita J. Serrano, Clara Lee, Robert J. Bryson-Richardson, Tamar E. Sztal

**Affiliations:** School of Biological Sciences, Monash University, Melbourne Australia

**Author notes:** Corresponding author Tamar E. Sztal.

**Keywords:** CDKL5 Deficiency Disorder, motor neurons, locomotion, microcephaly, zebrafish

## Abstract

Cyclin-dependent kinase-like-5 (CDKL5) Deficiency Disorder (CDD) is a severe X-linked neurodegenerative disease characterized by early-onset epileptic seizures, low muscle tone, progressive intellectual disability, severe motor function and visual impairment. CDD affects approximately 1 in 60,000 live births with many patients dying by early adulthood. For many patients, quality of life is significantly reduced due to the severity of their neurological symptoms and functional impairment. There are no effective therapies for CDD with current treatments focusing on improving symptoms rather than addressing the underlying causes of the disorder. Zebrafish offer a number of unique advantages for high-throughput pre-clinical evaluation of potential therapies for human neurological diseases including CDD. In particular, the large number of zebrafish that can be produced, together with the possibilities for *in vivo* imaging and genetic manipulation, allows for the detailed assessment of disease pathogenesis and therapeutic discovery. We have characterised a loss of function zebrafish model for CDD, containing a nonsense mutation in *cdkl5*. *cdkl5* mutant zebrafish display defects in neuronal patterning, microcephaly, and reduced muscle function caused by impaired muscle innervation. This study provides a powerful vertebrate model to investigate CDD disease pathophysiology and allow high-throughput screening for effective therapies.

## Introduction

CDKL5 deficiency disorder (CDD) is a severe neurodegenerative disease caused by mutations in *Cyclin-dependent kinase-like-5* (*CDKL5*). Patients with CDD often display a heterogeneous array of clinical phenotypes including early infantile epilepsy, delayed motor function and intellectual disability [32]. CDKL5 is a member of a highly conserved family of serine-threonine kinases, functioning to regulate cytoskeletal dynamics, synaptic vesicle stability and release, neurite outgrowth, and dendritic spine development [26]. To date, more than 265 variants have been identified in *CDKL5*, with approximately 70 of these considered pathogenic [15]. Many mutations are located within the catalytic domain, and those disrupting catalytic activity lead to a complete loss of kinase function [4, 15]. Due to the rarity of the disorder (1 in 60,000 live births) very little is known about long term prognosis, however, many patients die by early adulthood. For most patients, there are daily and ongoing medical challenges associated with managing the severe neurological impairments as well as the range of progressive and complex clinical comorbidities associated with the disease.

There are no effective therapies for CDD, with most treatments focusing on improving symptoms rather than addressing the underlying cause of the disorder. A handful of compounds are currently in clinical trials, including Ataleuren, currently used to treat Duchenne Muscular Dystrophy, to promote read-through of CDKL5 nonsense mutations [18] and ganaxolone to reduce seizure activity [37]. The lack of effective therapies is in part due to the complexity of the symptoms exhibited by patients and the limited understanding of CDKL5 function and its associated signal transduction pathways.

Previous studies have shown that Cdkl5 knockout (KO) mice recapitulate many features of CDD, exhibiting severe neuronal impairment leading to deficits in learning, memory and social behaviour [11, 23, 31, 34]. These deficiencies are associated with abnormalities in neurite growth and proliferation leading to reduced dendritic arborization of cortical neurons and spine density of hippocampal neurons affecting brain function and visual cortex processing [1, 34]. Significantly, neuronal development was shown to be arrested in Cdkl5 KO mice, suggesting that symptoms may be caused by improper dendritic maturation and formation of neuronal structures rather than degeneration of nervous tissues [5, 11, 12, 23]. However, the majority of Cdkl5 KO mice models display only minor motor function deficits with small alterations observed in their gait suggesting a mild coordination disturbance [1, 19, 34].

Unlike other vertebrates, the zebrafish model is highly suited to *in vivo* analysis of neurological development and CDD progression. Notably the optical clarity and power of live imaging, combined with accessibility of developing zebrafish larvae, provide novel ways to monitor disease states. Development of both the brain and spinal cord is highly conserved between humans and zebrafish, as well as many signalling pathways involved in their function. In zebrafish, the skeletal muscle makes up around 80% of the body, is functional as early as 20 hours post fertilization (hpf), and after 2 days post fertilization (dpf) the fish swim spontaneously. Muscle contractions controlling early spontaneous motor activity are likely to be directly modulated by embryonic central nervous system activity. At this time, the upper motor neurons in the primary cortex send their projections to the developing spinal cord and it is only after the fast muscle is formed at approximately 52 hpf, that the lower motor neurons project to the myotome [6].

Zebrafish are highly amenable to high-throughput pre-clinical evaluation of potential therapies for severe congenital disorders including CDD. Measurements of swimming behaviour using automated tracking systems can yield an unbiased, reliable, and high-throughput assessment of swimming performance that is reflective of skeletal muscle function and innervation [29].

In zebrafish, *cdkl5* exists as a single gene copy and bioinformatics analysis showed that Cdkl5 is highly conserved between zebrafish and its vertebrate orthologs, being 89% identical to the chicken, human, and mouse proteins [16]. Previous studies have reported two *cdkl5* transcripts (long and short) produced during zebrafish development, which differ in the inclusion of exon 16 [16]. Both isoforms are expressed during early zebrafish development, specifically in neural tissues including the brain and eye [16]. Mammals are known to have multiple splice variants, with the longest isoform also expressed highly in the human brain [9, 22].

Here we have characterized a zebrafish model for CDD to study disease progression and biology. We show that *cdkl5^−/−^* mutant zebrafish display reduced head size, defects in motor neuron branching leading to reduced muscle innervation, and impaired motor function. Using live confocal imaging, we show that *cdkl5^−/−^* mutant zebrafish also display microcephaly, a reduction in total brain volume and, in particular, a reduction in the cerebellum which regulates motor movements. Together our results validate this novel disease model which may be exploited for testing of potential therapeutics.

## Methods

### Zebrafish maintenance, morpholino injections and genotyping

Zebrafish were maintained according to standard protocols [35]. Zebrafish strains used were Tg(*islet1:eGFP*) [14], Tg(*HuC:eGFP*) [28], and an ENU-generated *cdkl5* mutant line (sa21938) obtained from the Zebrafish International Resource Centre. For *cdkl5* genotyping, allele specific PCR KASP technology (Geneworks) was used. For morpholino (MO) injections, the *cdkl5* ex5 (5’ AGATATAAACACTGTCATACCTCTG 3’) and Standard Control (5’ CCTCTTACCTCAGTTACAATTTATA 3’) MOs (GeneTools) were diluted in distilled water and co-injected with Cascade Blue labelled dextran (Molecular Probes) into one- to two-cell wild-type (*Tübingen*, TU strain) embryos. MO concentrations were calibrated according to [38] at the indicated concentrations (0.125 mM, 0.25 mM, and 0.5 mM for the *cdkl5* ex5 MO corresponding to 0.5, 1.0, or 2.0 ng; and 0.5 mM for the Standard Control MO corresponding to 2.0 ng). At 1 dpf, the embryos were sorted for Cascade Blue labelling.

### Whole-mount in situ hybridisation

Whole-mount *in situ* hybridization was carried out as described previously [25]. Probes were constructed using *cdkl5* specific primers F: GGATCTAAACGCAGCTCCTG and R: GGATCTAAACGCAGCTCCTG.

### cDNA synthesis and reverse transcription-PCR

For the qRT-PCR experiment, head and tails were separated from 3 dpf zebrafish, the progeny of a *cdkl5^+/−^* incross, for three independent biological replicates. The tails were used for genotyping and the heads were used for RNA extraction with 20 embryos per genotype per replicate. For morpholino experiments, whole embryos were analysed at either 3 or 6 dpf. Total RNA was extracted using TRIzol reagent (Invitrogen Life Technologies). cDNA was synthesized from 1μg of each RNA sample in a 20 μl reaction using Protoscript first strand cDNA synthesis kit (New England Biosciences), oligo(dT)20 and Random Primer Mix primers following the supplier’s instructions. Primers used for RT-PCR were *cdkl5*F: CTCCGTACGCTCAAACAAGAC, *cdkl5*R: TGAGCTCTCCCAAAATACACC, *βAct*F: GCATTGCTGACCGTATGCAG, *βAct*R: GATCCACATCTGCTGGAAGGTGG, *ef1α*F: CTGGAGGCCAGCTCAAACAT and *ef1α*R: ATCAAGAAGAGTAGTACCGCTAGCATTAC.

### Locomotion assays

Locomotion assays were performed on 6 dpf zebrafish as per [29]. An inactivity threshold of 1 mm/sec, detection threshold of 25 mm/sec and maximum burst threshold of 30 mm/sec were used. The total distance swum above inactivity threshold and below maximum burst threshold in a 10-min period were extracted using the ZebraLab software (Viewpoint Life Sciences).

### Antibody staining and confocal microscopy

Immunofluorescence was performed on 2 and 3 dpf zebrafish as per [30]. Antibodies used were anti-α-Actinin2 (Sigma clone A7811, 1:200), anti-α-myosin heavy chain antibody (DSHB, clone A4.1025), anti-α-acetylated tubulin antibody (Sigma T6793) and an AlexaFluorTM-labelled-488 secondary antibody (Molecular Probes, 1:200). Imaging was carried out on an LSM 710 confocal microscope (Zeiss) equipped with a 20X 1.0 numerical aperture water dipping objective and a 488 nm or 561 nm laser.

### Alcian Blue staining

*cdkl5^+/−^* fish were incrossed and their progeny were raised in embryo media (NaCl 5 mM, KCl 0.17mM, CaCl_2_ 0.33 mM, MgSO_4_ 0.33 mM in water) containing N-Phenylthiourea 200 μM (PTU, Sigma) from 12 hours to suppress melanocyte formation, changing medium every 24 hours. At 7 dpf, embryos were fixed in 4% paraformaldehyde and then left overnight in methanol at 4°C. Embryos were then rinsed in acid alcohol (70% EtOH: 1% HCl) and then stained overnight at room temperature in 0.1% Alcian Blue: 80% EtOH: 20% Acetic Acid. Embryos were then washed for 6 hours in acid alcohol and then rinsed in PBST before imaging.

### Brain image registration and analysis

*cdkl5^+/−^* fish were crossed to the Tg(*HuC:eGFP*) line and raised to adulthood. Tg*(HuC:eGFP);cdkl5^+/−^* fish were crossed to *cdkl5^+/−^* fish, and their progeny were raised in embryo media containing N-Phenylthiourea 200 μM (PTU, Sigma) from 12 hpf to suppress melanocyte formation, changing medium every 24 hours. Embryos were sorted for fluorescence at 2 dpf. At 4 or 5 dpf, fish were anesthetized using 0.0016% Tricaine methanesulfonate (Sigma) in embryo media, DNA was extracted from a small tail clipping and was used to genotype embryos (as described above). At 6 dpf, embryos were again anesthetised and set in 1% low melting point agarose in E3 containing tricaine in 0.8 mm fluorinated ethylene propylene (FEP) tubing (Bola). Images were taken using a Thorlabs confocal microscope with an Olympus 20x water dipping 1.0 NA objective, pinhole 25 μm, 2.005 μm/pixel, step size = 1 μm, averaging = 16 frames. Brain image registration and total brain and cerebellar volume analysis was performed as per [7, 13], combining the results from all cerebellar regions to determine total cerebellar volume.

### Motor neuron analyses

*cdkl5^+/−^* fish were crossed to the Tg*(islet1:eGFP)* line and raised to adulthood. Tg*(islet1:eGFP);cdkl5^+/−^* fish were crossed to *cdkl5^+/−^* fish, and their progeny were sorted for fluorescence at 2 dpf. At 3 dpf, fish were anesthetized using 0.0016% Tricaine methanesulfonate (Sigma) in embryo media, and DNA was extracted from a small tail clipping and was used to genotype embryos (as described above). At 6 dpf, embryos were again anesthetised and set in 1% low melting point agarose in E3 containing tricaine and imaged using an LSM 710 confocal microscope (Zeiss) equipped with a 20X 1.0 numerical aperture water dipping objective and a 488 nm laser. Images were taken from comparable regions along the spinal cord for each fish. The total number of GFP positive motor neuron cell bodies in Tg*(islet1:eGFP)* zebrafish were counted from a maximum projection image using the cell counter function in Fiji. The axonal density was determined by quantifying the surface of fluorescence above background. Each maximum projection Tg*(islet1:eGFP)* zebrafish image was cropped to remove any cell bodies. The background was normalised across images, and an upper threshold level of 64000 and lower threshold level of 9000 was used for each image. The images were converted to a mask image and the percentage area of GFP positive neuronal projections was determined using Fiji.

### Statistics

For all experiments, the investigators were blinded to genotype during data capture and once the analyses were completed, the genotypes of the fish were resolved. For qRT-PCR and swimming analyses all values were normalized to the average value of *cdkl5^+/+^* siblings. All statistical analyses were performed in GraphPad Prism 7 using a one-way ANOVA for qRT-PCR (**Figure 1c**) and *cdkl5^−/−^* mutant swimming assays (**Figure 2a&b**), one-tailed t-test for motor neuron analyses (**Figure 3b&c**), and one-tailed t-test for brain measurement (**Figure 5b&c**) analyses, Alcian Blue staining (**Figure 6c**) and Cdkl5 MO swimming assays (**Supplementary Figure 2c**). Data in **Figure 3c**, **Figure 4b** and Supplementary **Figure 2d** did not pass normality test and so a Mann-Whitney test was performed to determine significance. All other data were tested for normal distribution and passed using D’Agnostino and Perron’s test for Gaussian distribution. For brain measurements (**Figure 5**), we performed an outlier test using the ROUT method and identified one outlier (*cdkl5^+/+^* embryo 4 sample from replicate 4), which was excluded from the analysis.

**Figure 1:**
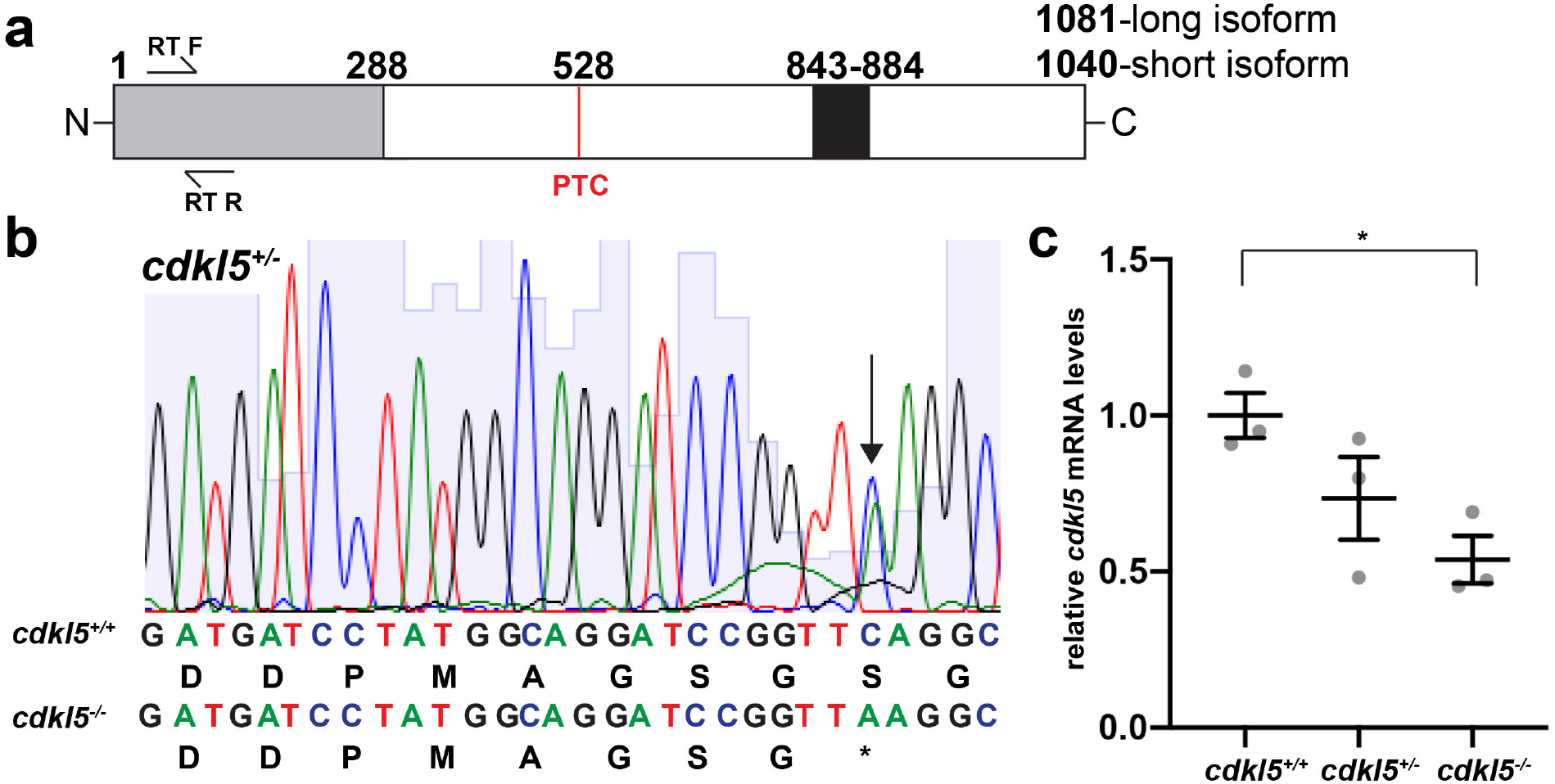
Validation of the *cdkl5^sa21938^* zebrafish mutant strain. a) Schematic of the zebrafish Cdkl5 protein showing the location of the nonsense (PTC) mutation, functional domains and primers used for quantitative RT-PCR analyses (grey = catalytic domain, black = ATP-binding region). b) *cdkl5^−/−^* mutants contain a C>A change in exon 11 of the gene (amino acid 528 out of 1040 and 1081 of the short and long isoforms respectively) creating a premature termination codon (PTC) as seen in the sequencing chromatograms from *cdkl5^+/−^* fish. c) Quantitative RT-PCR analysis shows that *cdkl5* mRNA levels are significantly reduced in *cdkl5^−/−^* embryos compared to *cdkl5^+/+^* siblings. Error bars represent mean±SEM range for three independent experiments (with 20 pooled fish per experiment), *p<0.05 using a one-tailed t-test.

**Figure 2:**
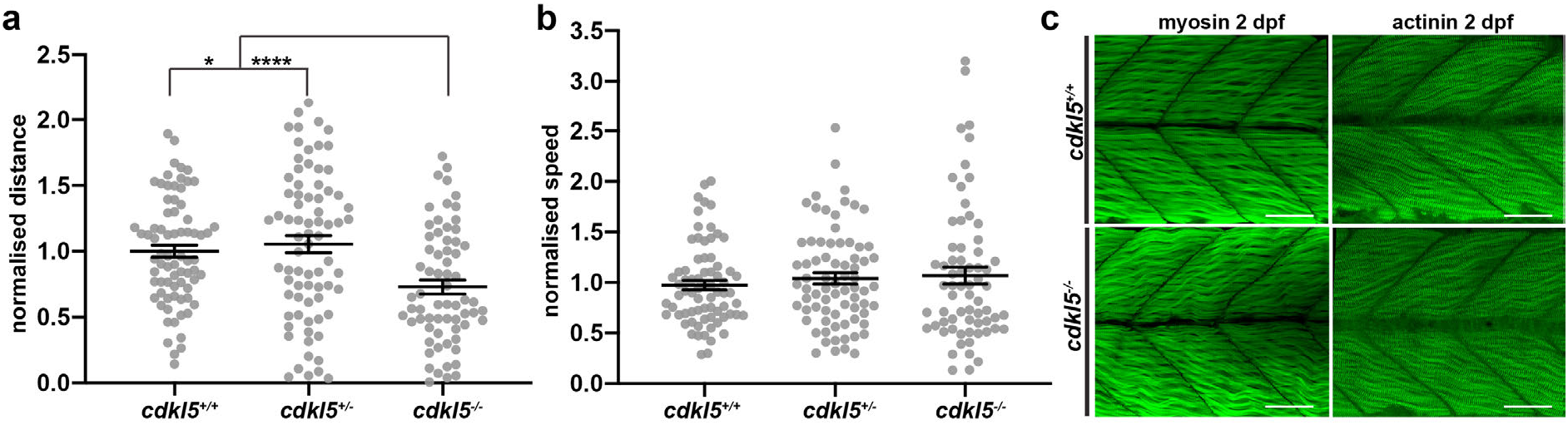
Motor function and muscle pathology of *cdkl5^−/−^* fish. a&b) Quantification of normalised a) distance travelled and b) speed of *cdkl5^+/+^*, *cdkl5^+/−^*, and *cdkl5^−/−^* fish at 6 dpf. Error bars represent mean±SEM for three independent experiments (n=38, 21, 15 *cdkl5^+/+^*and n=17, 39, 19 *cdkl5^+/−^* and n=27, 25, 14 *cdkl5^−/−^* fish per experiment), *p<0.05, **p<0.01 using a one-way ANOVA. c) Maximum projection of confocal image series of anti-Myosin and anti-Actinin2 antibody staining of *cdkl5^+/+^*and *cdkl5^−/−^* fish at 2 dpf. Scale bar = 50μm.

**Figure 3:**
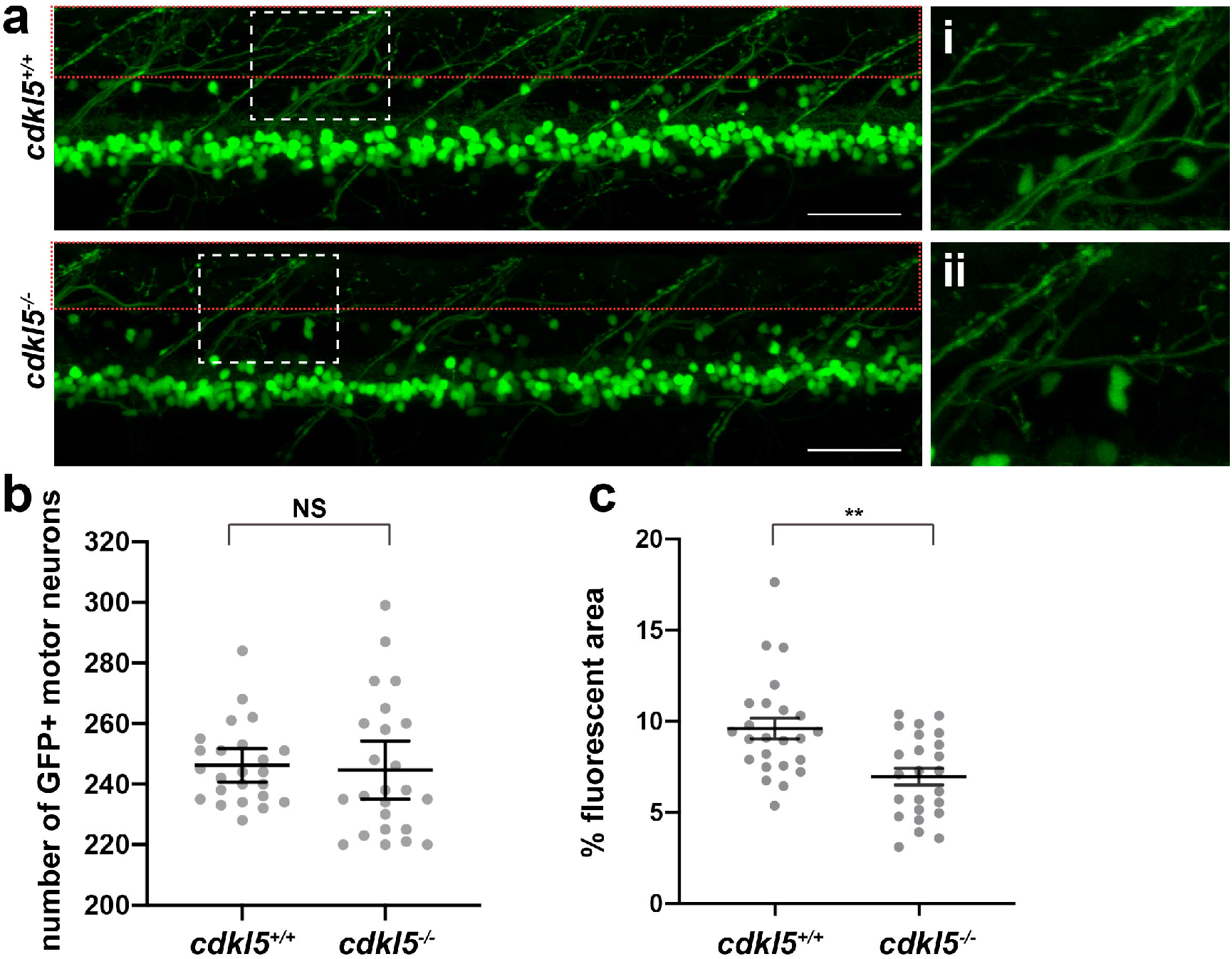
Assessment of neuronal pathology in *cdkl5^−/−^* mutants. a) Maximum projection confocal images of eGFP-labelled motor neurons from a) Tg(*islet1:eGFP); cdkl5^+/+^* (white dashed box zoomed in i) and Tg(*islet1:eGFP); cdkl5^−/−^* (white dashed box zoomed in ii) along the spinal region at 6 dpf. Red dashed box indicates the region quantified in 3c. b) Quantification of the number of GFP positive motor neurons at 6 dpf. Error bars represent mean±95% confidence interval for three independent experiments (n=8, 8, 8 Tg(*islet1:eGFP); cdkl5^+/+^* and n=8, 8, 8 Tg(*islet1:eGFP); cdkl5^−/−^* fish, NS = not significant, using a 2-tailed t-test. c) Quantification of the mean percent area of fluorescence from a) Tg(*islet1:eGFP); cdkl5^+/+^* and Tg(*islet1:eGFP); cdkl5^−/−^* (within the red dashed box region in 3a) at 6 dpf. Error bars represent mean±SEM range for three independent experiments (n=8, 8, 8 Tg(*islet1:eGFP); cdkl5^+/+^* and n=8, 8, 8 Tg(*islet1:eGFP); cdkl5^−/−^* fish, **p<0.01, using a one-tailed t-test. Scale bar = 50μm.

**Figure 4:**
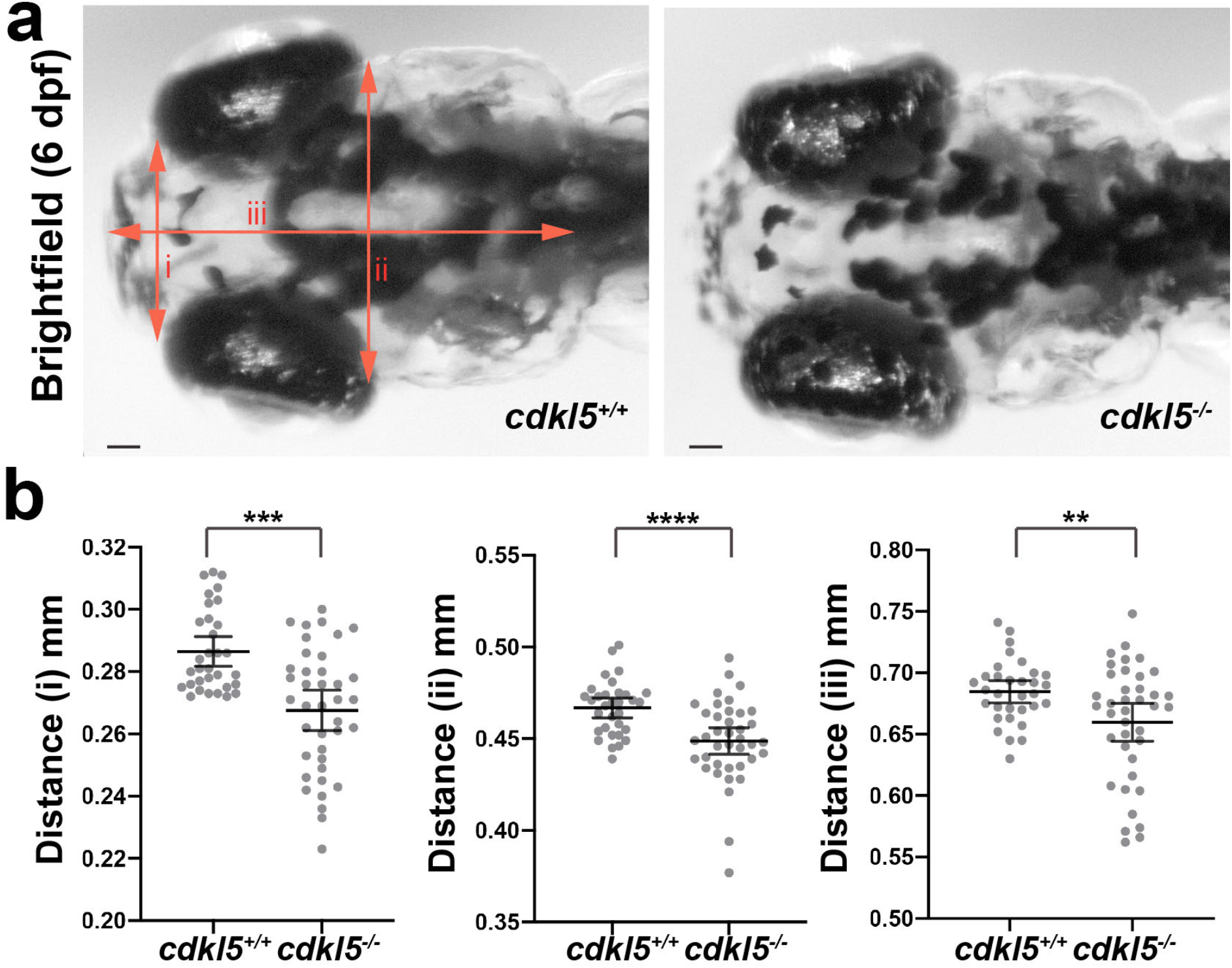
Analysis of head size in *cdkl5^−/−^* fish. a) Brightfield images of *cdkl5^−/−^* fish and *cdkl5^+/+^* sibling head regions at 6 dpf. Red lines (i: anterior head width, ii: posterior head width and iii: head length) indicate head region measured. b) Quantification of head regions (i: anterior head width, ii: posterior head width and iii: head length) of *cdkl5^−/−^* and *cdkl5^+/+^* fish. Error bars represent mean+95% confidence interval for four independent experiments (n=5, 8, 12, 7 for *cdkl5^+/+^* and n=8, 8, 10, 13 for *cdkl5^−/−^*), **p<0.01, ***p<0.001 and ****p<0.0001, using a one-tailed t-test. Scale bar = 50μm

**Figure 5:**
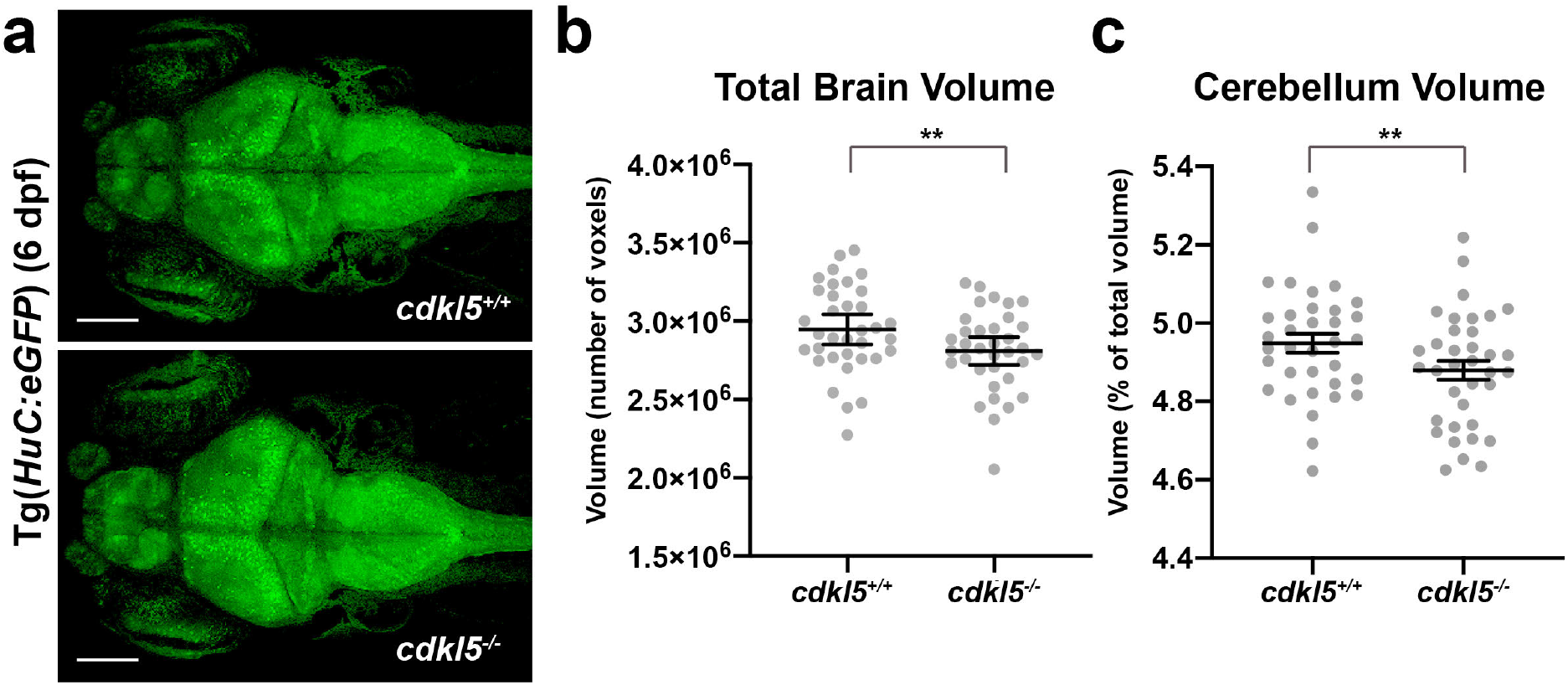
a) Maximum intensity projections of whole brains from Tg(*HuC:eGFP); cdkl5^+/+^* and Tg(*HuC:eGFP); cdkl5^−/−^* fish at 6 dpf. b&c) Quantification of b) total brain and c) cerebellum volume from *cdkl5^−/−^* and *cdkl5^+/+^* fish. Error bars represent mean±SEM for three independent experiments (n=12, 12, 11 *cdkl5^+/+^* and n=12, 12, 12 *cdkl5^−/−^* fish), **p<0.01, using a one-tailed t-test. Scale bar = 100μm.

**Figure 6:**
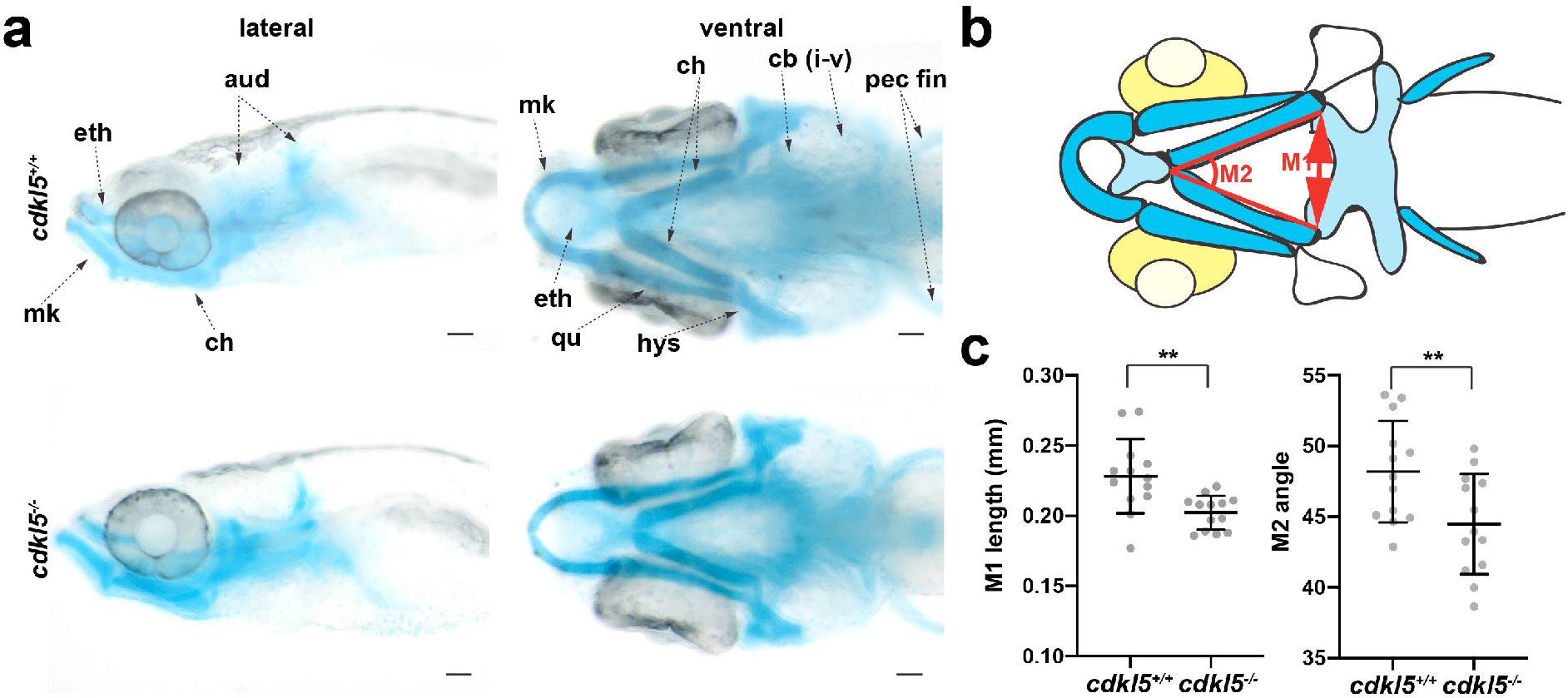
Alcian Blue staining reveals craniofacial cartilage structures in *cdkl5^−/−^* fish. a) Dorsal and ventral images of Alcian Blue staining in *cdkl5^−/−^* and *cdkl5^+/+^* fish at 7 dpf. b) Schematic of Alcian Blue staining in zebrafish heads at 7 dpf, showing locations of head measurements. (aud=auditory capsule, cb=ceratobranchials or gill arch derivatives, ch=ceratohyal, eth=ethmoid plate, hys=hyosymplectic, mk=Meckel’s cartilage, qu=quadrate, pec fin=pectoral fins) c) Quantification of the length (M1) and intersecting angle (M2) of the ceratohyal cartilage structures in *cdkl5^−/−^* and *cdkl5^+/+^* siblings at 7 dpf. Error bars represent mean+SD (n=13 fish per genotype), **p<0.01, using a one-tailed t-test. Scale bar = 100μm.

## Results

### cdkl5 is expressed during early zebrafish development

Consistent with previous reports [16, 33], we observed that *cdkl5* is expressed in the developing embryo (**Supplementary Figure 1a**), suggesting it plays a role in early zebrafish development. To determine the spatial expression of *cdkl5*, we performed *in situ* hybridization and found that *cdkl5* is expressed in the brain from 1 dpf, in the muscle at 2 dpf, in the otic vesicles, eyes and heart from 2 dpf, in the kidneys from 3 dpf and in the developing spinal cord from 4 dpf (**Supplementary Figure 1b**). The brain, spinal cord, eyes, muscle, and kidneys represent tissues that are affected in patients carrying *CDKL5* mutations [15] indicating that zebrafish may be a suitable model to study CDD pathogenesis.

### Loss of cdkl5 result in impaired muscle movement and neuronal branching

We obtained a *cdkl5* zebrafish mutant containing a C>A change in exon 11 of the gene (amino acid 528 of both the short and long isoforms) creating a premature termination codon leading to a truncated and non-functional gene product. We verified the mutation in this strain by Sanger sequencing and its location is predicted to affect both the long and short Cdkl5 isoforms (**Figure 1a&b**). Using quantitative RT-PCR, we determined that the mutation leads to a significant reduction in *cdkl5* mRNA in *cdkl5^−/−^* fish compared to their wild-type siblings (**Figure 1c**).

Patients carrying *CDLK5* mutations display motor function impairment, therefore we tested the swimming ability of *cdkl5* mutant zebrafish to determine whether motor function was affected. We detected a significant decrease in the normalised distance swum by *cdkl5^−/−^* fish (0.729+0.0536 SEM) compared to both *cdkl5^+/−^* (1.055+0.0647 SEM) and *cdkl5^+/+^* siblings (1.00+0.0469 SEM) (**Figure 2a**). Interestingly there was no significant difference in the swimming speed of *cdkl5^−/−^* fish (1.068+0.083 SEM) compared to their siblings (0.947+0.047 SEM for *cdkl5^+/+^* and 1.041+0.055 SEM for *cdkl5^+/−^*) (**Figure 2b**). We analysed the muscle pathology by labelling the trunk muscle for Myosin and Actinin2, essential components of the muscle fibre. We observed no overt difference in either muscle structure or patterning in *cdkl5^−/−^* mutants compared to wild-type siblings (**Figure 2c**) at 2 dpf.

We then crossed *cdkl5^+/−^* fish to a Tg*(islet1:eGFP)* zebrafish transgenic strain, labelling all of the motor neurons with enhanced green fluorescent protein (GFP) [14]. Using this transgenic approach, we were able to image motor neurons in the live embryo at 6 dpf to determine whether there were significant deviations in cell number and axon formation. Whilst we recorded no significant difference in the number of cell bodies in Tg*(islet1:eGFP)*;*cdkl5^−/−^* mutants (244.6±2.65 SEM) compared to Tg*(islet1:eGFP)*;*cdkl5^+/+^* siblings (246.2±5.96 SEM) (**Figure 3a&b**), we observed a significant reduction in the density of neuronal processes emerging from the spinal cord revealed by a reduction of fluorescent surface area in Tg*(islet1:eGFP)*;*cdkl5^−/−^* mutants (7.004±0.626 SEM) compared to Tg*(islet1:eGFP)*;*cdkl5^+/+^* siblings (9.600±0.499 SEM) (**Figure 3a&c**). This reduced axonal branching likely results in dysconnectivity between the spinal cord and adjacent muscle fibres.

### Morpholino knockdown of Cdkl5 recapitulated phenotypes observed in cdkl5 mutants

To confirm that a reduction in Cdkl5 causes defects in motor function and muscle innervation, we used a MO targeting the splice donor site of exon 5 (*cdkl5* ex5 MO) to knockdown Cdkl5. To determine an effective dose for knockdown, we injected the MO at 0.5 ng, 1.0 ng, or 2.0 ng into wild-type embryos and performed RT-PCR. We were able to detect a number of smaller sized amplicons by RT-PCR only in the 2.0 ng injected samples (**Supplementary Figure 2a**). Based on these observations, we selected 2.0 ng MO to use in further experiments. To confirm that *cdkl5* mRNA levels are reduced in Cdkl5 morphants at 6 dpf, we performed RT-PCR analyses and found that indeed *cdkl5* mRNA levels are decreased compared to Control MO-injected fish (**Supplementary Figure 2b**). We performed locomotion assays at 6 dpf and found a significant decrease in swimming performance in Cdkl5 morphants (0.683±0.031 SEM) compared to Control MO-injected fish (0.980±0.061 SEM) (**Supplementary Figure 2c&d**).

To investigate neuronal pathologies in Cdkl5 morphants, we used an anti-α-acetylated tubulin antibody to stain mature neurons, including their axons. We observed no obvious differences in neuronal morphology or projections at 3 dpf between Cdkl5 knockdown and Control MO-injected fish (**Supplementary Figure 2e**). However, in Tg*(islet1:eGFP)*;Cdkl5 knockdown fish, there appears to be a reduction in the density of axonal projections (**Supplementary Figure 2f**), recapitulating the pathologies observed in *cdkl5^−/−^* mutant fish (**Figure 3b**) and confirming that reduction of Cdkl5 affects zebrafish neuromuscular connectivity.

### cdkl5 mutants display skeletal abnormalities and microcephaly

CDD patients were reported to exhibit a small head size with some demonstrating cortical atrophy identified through brain imaging [36]. We therefore measured the size of the zebrafish head by taking width and length measurements and observed a clear reduction in both the width and length of the head in *cdkl5^−/−^* fish (anterior head width: 0.263±0.0169 SD, posterior head width: 0.447±0.009 SD and head length: 0.654±0.034 SD) compared to *cdkl5^+/+^* siblings (anterior head width : 0.2880±0.011 SD, posterior head width: 0.465±0.009 SD and head length: 0.684±0.012 SD) at 6 dpf (**Figure 4a&b**), recapitulating microcephaly phenotypes observed in CDD patients. We also measured brain volume in *cdkl5^−/−^* zebrafish at 6 dpf using live confocal microscopy and a pan-neuronal fluorescent reporter strain (Tg*(HuC:eGFP*)) [28]. We observed a 5% reduction (p<0.01) in total brain volume in Tg*(HuC:eGFP);cdkl5^−/−^* fish (2811034+30820 SEM) compared to Tg*(HuC:eGFP);cdkl5^+/+^* wild-type siblings (2944860+26472 SEM) (**Figure 5a&b**), in line with our previous observations (**Figure 4b**). We also measured the total volume of the cerebellum, responsible for coordinating voluntary movements (comprising the Eminentia granularis, Valvula cerebelli, Corpus cerebelli, and Cerebellar neuropil). We recorded a 1.4% reduction (p<0.01) in the cerebellum of Tg*(HuC:eGFP);cdkl5^−/−^* fish (4.880±0.0153 SEM) compared to Tg*(HuC:eGFP);cdkl5^+/+^* wild-type siblings (4.948±0.0162 SEM) (**Figure 5c**).

The cranial cartilage of the zebrafish is the first skeletal structure present and forms at approximately 3 dpf. As zebrafish grow, the bones continue to ossify, with the first osteoblasts surrounding the cartilage and forming bone matrix by 7 dpf [3]. We stained the developing cartilage using an Alcian Blue stain (**Figure 6a**) and found both a decrease in both the distance and angle between the two ceratohyal cartilage elements in *cdkl5^−/−^* fish (distance: 0.202+0.012 SD and angle: 44.47+0.9822) compared to *cdkl5^+/+^* siblings (distance: 0.228+0.026 SD and angle: 48.18+0.996) at 7 dpf (**Figure 6b&c**). This provides further evidence that *cdkl5^−/−^* fish display microcephaly.

## Discussion and conclusions

To date, no effective therapies have been identified for CDD with current treatments focusing on the alleviation of symptoms rather than addressing the underlying biology of the disease. A vertebrate animal model that can reliably be used to identify effective therapies is therefore needed. Here we describe the characterisation of a novel zebrafish model for CDD. *cdkl5^−/−^* mutant zebrafish display reduced muscle function, impaired neuronal formation, and microcephaly consistent with phenotypes observed in CDD patients.

So far, more than 265 variants have been reported in *CDKL5*, with mutations distributed all along the length of the protein [15]. Approximately 27% of these are considered pathogenic with many of these located in the catalytic domain [15]. Studies have shown that patients with mutations within the catalytic domain and frameshift mutations located at the end of the C-terminal region had more severe motor impairment, refractive (or non-drug responsive) epilepsy, or microcephaly [27]. Milder forms of the disease were caused by mutations in the ATP-binding region or nonsense mutations in the C-terminal regions with patients possessing better hand coordination and the ability to walk unaided [2, 20]. The nonsense mutation in *cdkl5^−/−^* mutant zebrafish sits 240 amino acids downstream of the catalytic domain and activates nonsense mediated decay pathways causing degradation of the resultant mutant mRNA. Therefore, we expect that phenotypes observed in *cdkl5^−/−^* mutant zebrafish will be comparable in severity to those contained within the catalytic domain.

Despite the extensive characterisation of neurological phenotypes, there is still not a clear correlation between type of mutation and phenotypic severity, however, delayed neurodevelopment appears to be a hallmark feature of CDD [2]. CDKL5 belongs to the serine-threonine kinase family and is an important factor influencing neuronal functions [17]. Interestingly, the levels of CDKL5 vary at different stages of development, with lowest expression in the prenatal stages and increasing throughout postnatal development [2, 26]. The changes in the levels of CDKL5 are consistent with a role in the process of neuronal formation, growth, and, in particular, in dendrite development and bifurcation [15]. We found that neuromuscular connectivity is impaired in *cdkl5^−/−^* zebrafish during early embryonic stages. Interestingly, we observed no significant difference in the number of motor neurons along the spinal cord but recorded a marked decrease in the density of motor neuron processes emerging from the spinal cord in *cdkl5^−/−^* mutants at 6 dpf, consistent with phenotypes observed in CDD patients and CDKL5 KO mice [11, 21]. At this age, *cdkl5^−/−^* mutant zebrafish displayed reduced swimming however, we did not detect any defects in muscle formation or patterning during early muscle development. Defects in motor neuron axon length and branching have been shown, in zebrafish models of Motor Neuron Disease, to correlate with shorter swimming distances [24]. This supports the idea that the defects in swimming performance of *cdkl5^−/−^* mutant zebrafish may be caused by impaired motor neuron outputs.

We have also shown that *cdkl5^−/−^* mutant zebrafish displayed reduced head size and a reduction in brain volume [36]. In CDD patients, head circumference at birth is normal in approximately 93% of cases however, in the first few years of life, head circumference is reduced [10]. In severe cases reduced head size and microcephaly is often accompanied by encephalopathy causing profound mental retardation [8]. Examination of the cerebellum revealed a clear reduction in cerebellum volume in *cdkl5^−/−^* animals compared to their wild-type siblings. The cerebellum is the major brain region responsible for controlling voluntary movements including motor function, balance, and coordination, and defects in cerebellum formation may contribute to impaired motor function in *cdkl5^−/−^* fish. In conclusion, our study has validated a novel zebrafish model for CDD that will provide mechanistic insight into CDD biology and is also an excellent system in which to perform high-throughput chemical screening for effective therapies.

## Declarations

### Ethics statement

Fish maintenance and handling was carried out as per the standard operating procedures approved by the Monash Animal Services Animal Ethics Committee under breeding colony license MARP/2015/004/BC. Fish were anaesthetized using Tricaine methanesulfonate.

### Consent for publication

Not applicable.

### Availability of data and materials

The data is made available on Bridges using the following link: https://figshare.com/projects/Novel_pre-clinical_model_for_CDKL5_Deficiency_Disorder/91460

### Authors’ contribution

TS performed the experiments and analysed the data as well as wrote, reviewed and edited the manuscript. RS, CL, and RBR performed the brain imaging and alignment, RS, CL, and RBR edited and reviewed the manuscript.

## Acknowledgements

We would like to thank Dr Jihane Homman-Ludiye for critically reading and providing feedback on the manuscript. The myosin antibody (clone A4.1025) was obtained from the Developmental Studies Hybridoma Bank, created by the NICHD of the NIH and maintained at The University of Iowa, Department of Biology, Iowa City, IA 52242.

## References

1 Amendola E, Zhan Y, Mattucci C, Castroflorio E, Calcagno E, Fuchs C, Lonetti G, Silingardi D, Vyssotski AL, Farley Det al (2014) Mapping pathological phenotypes in a mouse model of CDKL5 disorder. PLoS One 9: e91613 Doi 10.1371/journal.pone.0091613

2 Bahi-Buisson N, Bienvenu T (2012) CDKL5-Related Disorders: From Clinical Description to Molecular Genetics. Molecular Syndromology 2: 137–152 Doi 10.1159/000331333

3 Bergen DJM, Kague E, Hammond CL (2019) Zebrafish as an Emerging Model for Osteoporosis: A Primary Testing Platform for Screening New Osteo-Active Compounds. Frontiers in Endocrinology 10: 6 Doi 10.3389/fendo.2019.00006

4 Bertani I, Rusconi L, Bolognese F, Forlani G, Conca B, De Monte L, Badaracco G, Landsberger N, Kilstrup-Nielsen C (2006) Functional consequences of mutations in CDKL5, an X-linked gene involved in infantile spasms and mental retardation. The Journal of Biological Chemistry 281: 32048–32056 Doi 10.1074/jbc.M606325200

5 Chen Q, Zhu Y-C, Yu J, Miao S, Zheng J, Xu L, Zhou Y, Li D, Zhang C, Tao Jet al (2010) CDKL5, a Protein Associated with Rett Syndrome, Regulates Neuronal Morphogenesis via Rac1 Signaling. The Journal of Neuroscience 30: 12777–12786 Doi 10.1523/jneurosci.1102-10.2010

6 D’Elia KP, Dasen JS (2018) Development, functional organization, and evolution of vertebrate axial motor circuits. Neural Development 13: 10 Doi 10.1186/s13064-018-0108-7

7 Dark C, Williams C, Bellgrove MA, Hawi Z, Bryson-Richardson RJ (2020) Functional validation of CHMP7 as an ADHD risk gene. Translational psychiatry 10: 385 Doi 10.1038/s41398-020-01077-w

8 Fehr S, Downs J, Ho G, de Klerk N, Forbes D, Christodoulou J, Williams S, Leonard H (2016) Functional abilities in children and adults with the CDKL5 disorder. American Journal of Medical Genetics Part A 170: 2860–2869 Doi 10.1002/ajmg.a.37851

9 Fichou Y, Nectoux J, Bahi-Buisson N, Chelly J, Bienvenu T (2011) An isoform of the severe encephalopathy-related *CDKL5* gene, including a novel exon with extremely high sequence conservation, is specifically expressed in brain. Journal of Human Genetics 56: 52–57 Doi 10.1038/jhg.2010.143

10 Frullanti E, Papa FT, Grillo E, Clarke A, Ben-Zeev B, Pineda M, Bahi-Buisson N, Bienvenu T, Armstrong J, Roche Martinez Aet al (2019) Analysis of the Phenotypes in the Rett Networked Database. International Journal of Genomics 2019: 6956934 Doi 10.1155/2019/6956934

11 Fuchs C, Gennaccaro L, Trazzi S, Bastianini S, Bettini S, Lo Martire V, Ren E, Medici G, Zoccoli G, Rimondini Ret al (2018) Heterozygous CDKL5 Knockout Female Mice Are a Valuable Animal Model for CDKL5 Disorder. Neural Plasticity 2018: 9726950 Doi 10.1155/2018/9726950

12 Fuchs C, Trazzi S, Torricella R, Viggiano R, De Franceschi M, Amendola E, Gross C, Calzà L, Bartesaghi R, Ciani E (2014) Loss of CDKL5 impairs survival and dendritic growth of newborn neurons by altering AKT/GSK-3β signaling. Neurobiology of Disease 70: 53–68 Doi 10.1016/j.nbd.2014.06.006

13 Gupta T, Marquart GD, Horstick EJ, Tabor KM, Pajevic S, Burgess HA (2018) Morphometric analysis and neuroanatomical mapping of the zebrafish brain. Methods (San Diego, Calif) 150: 49–62 Doi 10.1016/j.ymeth.2018.06.008

14 Higashijima S, Hotta Y, Okamoto H (2000) Visualization of cranial motor neurons in live transgenic zebrafish expressing green fluorescent protein under the control of the islet-1 promoter/enhancer. The Journal of Neuroscience 20: 206–218 Doi 10.1523/jneurosci.20-01-00206.2000

15 Jakimiec M, Paprocka J, Śmigiel R (2020) CDKL5 Deficiency Disorder-A Complex Epileptic Encephalopathy. Brain sciences 10: Doi 10.3390/brainsci10020107

16 Katayama S, Senga Y, Oi A, Miki Y, Sugiyama Y, Sueyoshi N, Kameshita I (2016) Expression analyses of splice variants of zebrafish cyclin-dependent kinase-like 5 and its substrate, amphiphysin 1. Gene 583: 15–23 Doi 10.1016/j.gene.2016.02.036

17 Kilstrup-Nielsen C, Rusconi L, La Montanara P, Ciceri D, Bergo A, Bedogni F, Landsberger N (2012) What we know and would like to know about CDKL5 and its involvement in epileptic encephalopathy. Neural Plasticity 2012: 728267 Doi 10.1155/2012/728267

18 McDonald CM, Campbell C, Torricelli RE, Finkel RS, Flanigan KM, Goemans N, Heydemann P, Kaminska A, Kirschner J, Muntoni Fet al (2017) Ataluren in patients with nonsense mutation Duchenne muscular dystrophy (ACT DMD): a multicentre, randomised, double-blind, placebo-controlled, phase 3 trial. Lancet (London, England) 390: 1489–1498 Doi 10.1016/s0140-6736(17)31611-2

19 Okuda K, Takao K, Watanabe A, Miyakawa T, Mizuguchi M, Tanaka T (2018) Comprehensive behavioral analysis of the Cdkl5 knockout mice revealed significant enhancement in anxiety- and fear-related behaviors and impairment in both acquisition and long-term retention of spatial reference memory. PLoS One 13: e0196587 Doi 10.1371/journal.pone.0196587

20 Olson HE, Demarest ST, Pestana-Knight EM, Swanson LC, Iqbal S, Lal D, Leonard H, Cross JH, Devinsky O, Benke TA (2019) Cyclin-Dependent Kinase-Like 5 Deficiency Disorder: Clinical Review. Pediatric Neurology 97: 18–25 Doi 10.1016/j.pediatrneurol.2019.02.015

21 Pizzo R, Gurgone A, Castroflorio E, Amendola E, Gross C, Sassoè-Pognetto M, Giustetto M (2016) Lack of Cdkl5 Disrupts the Organization of Excitatory and Inhibitory Synapses and Parvalbumin Interneurons in the Primary Visual Cortex. Frontiers in Cellular Neuroscience 10: 261 Doi 10.3389/fncel.2016.00261

22 Rademacher N, Hambrock M, Fischer U, Moser B, Ceulemans B, Lieb W, Boor R, Stefanova I, Gillessen-Kaesbach G, Runge Cet al (2011) Identification of a novel *CDKL5* exon and pathogenic mutations in patients with severe mental retardation, early-onset seizures and Rett-like features. Neurogenetics 12: 165–167 Doi 10.1007/s10048-011-0277-6

23 Ren E, Roncacé V, Trazzi S, Fuchs C, Medici G, Gennaccaro L, Loi M, Galvani G, Ye K, Rimondini Ret al (2019) Functional and Structural Impairments in the Perirhinal Cortex of a Mouse Model of CDKL5 Deficiency Disorder Are Rescued by a TrkB Agonist. Frontiers in Cellular Neuroscience 13: 169 Doi 10.3389/fncel.2019.00169

24 Robinson KJ, Yuan KC, Don EK, Hogan AL, Winnick CG, Tym MC, Lucas CW, Shahheydari H, Watchon M, Blair IPet al (2019) Motor Neuron Abnormalities Correlate with Impaired Movement in Zebrafish that Express Mutant Superoxide Dismutase 1. Zebrafish 16: 8–14 Doi 10.1089/zeb.2018.1588

25 Ruparelia AA, Zhao M, Currie PD, Bryson-Richardson RJ (2012) Characterization and investigation of zebrafish models of filamin-related myofibrillar myopathy. Hum Mol Genet 21: 4073–4083 Doi 10.1093/hmg/dds231

26 Rusconi L, Salvatoni L, Giudici L, Bertani I, Kilstrup-Nielsen C, Broccoli V, Landsberger N (2008) CDKL5 expression is modulated during neuronal development and its subcellular distribution is tightly regulated by the C-terminal tail. The Journal of Biological Chemistry 283: 30101–30111 Doi 10.1074/jbc.M804613200

27 Russo S, Marchi M, Cogliati F, Bonati MT, Pintaudi M, Veneselli E, Saletti V, Balestrini M, Ben-Zeev B, Larizza L (2009) Novel mutations in the *CDKL5* gene, predicted effects and associated phenotypes. Neurogenetics 10: 241–250 Doi 10.1007/s10048-009-0177-1

28 St John JA, Key B (2012) HuC-eGFP mosaic labelling of neurons in zebrafish enables *in vivo* live cell imaging of growth cones. Journal of Molecular Histology 43: 615–623 Doi 10.1007/s10735-012-9462-7

29 Sztal TE, Ruparelia AA, Williams C, Bryson-Richardson RJ (2016) Using Touch-evoked Response and Locomotion Assays to Assess Muscle Performance and Function in Zebrafish. Journal of Visualized Experiments : JoVE: Doi 10.3791/54431

30 Sztal TE, Zhao M, Williams C, Oorschot V, Parslow AC, Giousoh A, Yuen M, Hall TE, Costin A, Ramm Get al (2015) Zebrafish models for nemaline myopathy reveal a spectrum of nemaline bodies contributing to reduced muscle function. Acta Neuropathologica 130: 389–406 Doi 10.1007/s00401-015-1430-3

31 Tang S, Wang IJ, Yue C, Takano H, Terzic B, Pance K, Lee JY, Cui Y, Coulter DA, Zhou Z (2017) Loss of CDKL5 in Glutamatergic Neurons Disrupts Hippocampal Microcircuitry and Leads to Memory Impairment in Mice. The Journal of Neuroscience 37: 7420–7437 Doi 10.1523/jneurosci.0539-17.2017

32 Tao J, Van Esch H, Hagedorn-Greiwe M, Hoffmann K, Moser B, Raynaud M, Sperner J, Fryns JP, Schwinger E, Gécz Jet al (2004) Mutations in the X-linked cyclin-dependent kinase-like 5 (*CDKL5/STK9*) gene are associated with severe neurodevelopmental retardation. American Journal of Human Genetics 75: 1149–1154 Doi 10.1086/426460

33 Vitorino M, Cunha N, Conceição N, Cancela ML (2018) Expression pattern of cdkl5 during zebrafish early development: implications for use as model for atypical Rett syndrome. Molecular Biology Reports 45: 445–451 Doi 10.1007/s11033-018-4180-1

34 Wang IT, Allen M, Goffin D, Zhu X, Fairless AH, Brodkin ES, Siegel SJ, Marsh ED, Blendy JA, Zhou Z (2012) Loss of CDKL5 disrupts kinome profile and event-related potentials leading to autistic-like phenotypes in mice. Proc Natl Acad Sci U S A 109: 21516–21521 Doi 10.1073/pnas.1216988110

35 Westerfield M (2007) The Zebrafish Book. A guide for the laboratory use of zebrafish (Danio rerio). Univ of Oregon Press, City

36 Yamamoto T, Shimojima K, Kimura N, Mogami Y, Usui D, Takayama R, Ikeda H, Imai K (2015) Recurrent occurrences of *CDKL5* mutations in patients with epileptic encephalopathy. Human Genome Variation 2: 15042 Doi 10.1038/hgv.2015.42

37 Yawno T, Miller SL, Bennet L, Wong F, Hirst JJ, Fahey M, Walker DW (2017) Ganaxolone: A New Treatment for Neonatal Seizures. Frontiers in cellular Neuroscience 11: 246 Doi 10.3389/fncel.2017.00246

38 Yuan S, Sun Z (2009) Microinjection of mRNA and morpholino antisense oligonucleotides in zebrafish embryos. Journal of Visualized Experiments : JoVE: Doi 10.3791/1113

