## Supplementary Figures for "Novel pre-clinical model for CDKL5 Deficiency Disorder"

Supplementary material

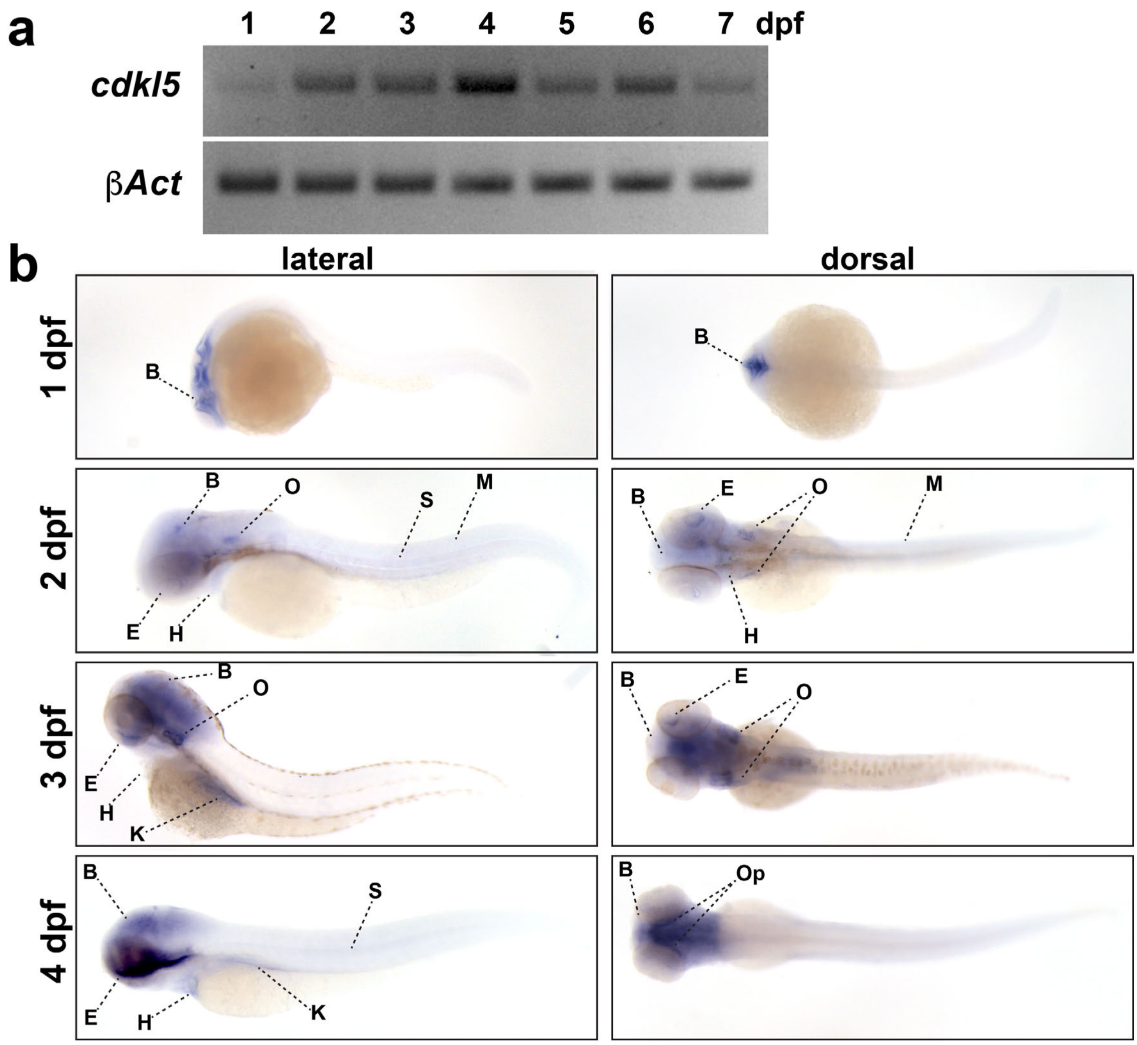


**Supplementary Figure 1:** *cdkl5* expression during early zebrafish development. a) RT-PCR analyses of *cdkl5* from 1-7 dpf. *β-Act* was amplified as a positive control. b) *in situ* hybridisation of *cdkl5* in 1-4 dpf zebrafish. At 1 dpf, *cdkl5* is detected in the brain and at low levels in the skeletal muscle (along the trunk of the fish). At 2 dpf, *cdkl5* mRNA is detected in the brain, heart, eyes, otic (or auditory) vesicles, and at low levels in the muscle. At 3 dpf, *cdkl5* is detected in the brain, heart, eyes, and otic vesicles. At 4 dpf, *cdkl5* is detected in the brain regions, heart, eyes, and developing spinal cord. (B=brain, E=eye, H=heart, K=kidney, O=otic vesicles, Op=optic tectum, M=muscle, S=spinal cord).

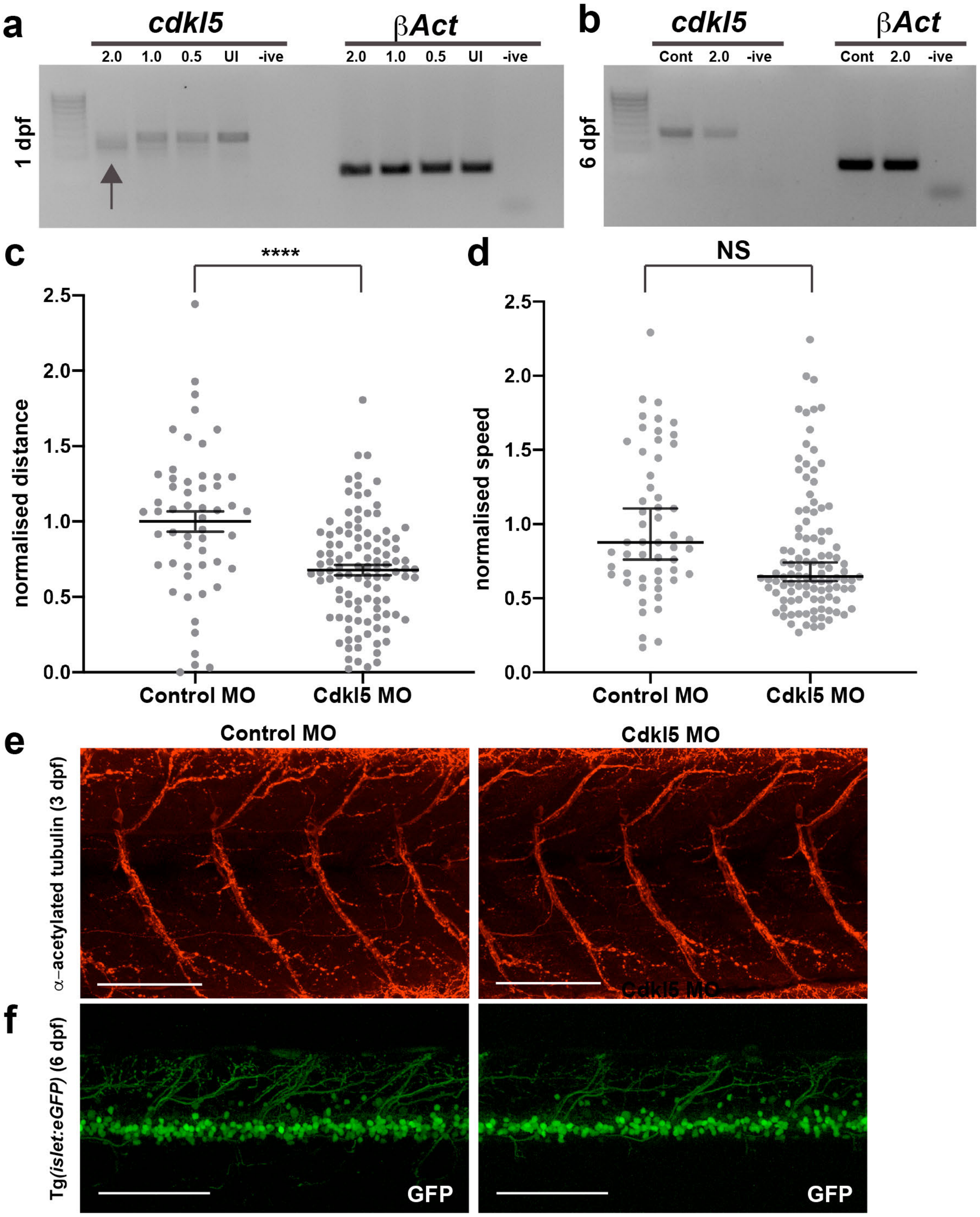
a
cdkl5
βAct
2.0 1.0 0.5 UI -ive
2.0 1.0 0.5 UI -ive
1 dpf
b
cdkl5
βAct
Cont 2.0 -ive
Cont 2.0 -ive
6 dpf
c
****
normalised distance
2.5
2.0
1.5
1.0
0.5
0.0
Control MO
Cdkl5 MO
d
NS
normalised speed
2.5
2.0
1.5
1.0
0.5
0.0
Control MO
Cdkl5 MO
e
Control MO
Cdkl5 MO
α-acetylated tubulin (3 dpf)
f
Tg(islet:eGFP) (6 dpf)
GFP
GFP

**Supplementary Figure 2:** Morpholino knockdown of Cdkl5. a) RT-PCR analysis for *cdkl5* mRNA following Cdkl5 MO injection. The amplicon in uninjected fish is the expected product size of 424 bp and is present with decreasing intensity in the Cdkl5 MO-injected fish as the injected concentration of MO increases from 0.5 ng to 2.0 ng. The lower bands (arrow) appear in the Cdkl5 MO-injected fish at the 2.0 ng concentration and results from mis-splicing of the *cdkl5* mRNA. b) At 6 dpf, the correct band is present in Cdkl5 MO-injected fish, however, it is diminished in intensity, demonstrating a reduction in *cdkl5* mRNA levels compared to Control MO-injected fish. c&d) Quantification of normalised c) distance travelled and d) speed of Cdkl5 MO-injected and Control MO-injected fish at 6 dpf. Error bars for distance and speed represent mean$\pm$SEM and median$\pm$interquartile range respectively for three independent experiments (n=18, 17, 19 Control MO-injected fish and n=27, 39, 40 Cdkl5 MO-injected fish), ***$p<0.001$, NS=not significant, using a one-way ANOVA. e) Maximum projection confocal images of $\alpha$-acetylated tubulin antibody staining of Cdkl5 MO and Control MO-injected fish fish at 3 dpf. f) GFP-labelled motor neurons in the spinal cord of Tg(*islet1:eGFP*) injected with either the Cdkl5 MO or Control MO at 6 dpf. A reduction in neuronal projections is evident. Scale bar = 100μm.
